# Functional Organization of Auditory and Reward Systems in Aging

**DOI:** 10.1101/2023.01.01.522417

**Authors:** Alexander Belden, Milena Aiello Quinci, Maiya Geddes, Nancy J. Donovan, Suzanne B. Hanser, Psyche Loui

## Abstract

The intrinsic organization of functional brain networks is known to change with age, and is affected by perceptual input and task conditions. Here, we compare functional activity and connectivity during music listening and rest between younger (N=24) and older (N=24) adults, using whole brain regression, seed-based connectivity, and ROI-ROI connectivity analyses. As expected, activity and connectivity of auditory and reward networks scaled with liking during music listening in both groups. Younger adults show higher within-network connectivity of auditory and reward regions as compared to older adults, both at rest and during music listening, but this age-related difference at rest was reduced during music listening, especially in individuals who self-report high musical reward. Furthermore, younger adults showed higher functional connectivity between auditory network and medial prefrontal cortex (mPFC) that was specific to music listening, whereas older adults showed a more globally diffuse pattern of connectivity, including higher connectivity between auditory regions and bilateral lingual and inferior frontal gyri. Finally, connectivity between auditory and reward regions was higher when listening to music selected by the participant. These results highlight the roles of aging and reward sensitivity on auditory and reward networks. Results may inform the design of music-based interventions for older adults, and improve our understanding of functional network dynamics of the brain at rest and during a cognitively engaging task.

## Introduction

The process of aging is characterized by changes in a variety of brain functions that underlie motivated behavior (Mather, 2016). One set of age-related changes involves a loss of connectivity in functional network dynamics of the brain (Tomasi & Volkow, 2012; Sala-Llonch, Bartrés-Faz & Junqué, 2015; Grady et al, 2016). The default mode network (DMN), for example, shows decreased connectivity in normal aging (Hafkemeijer et al, 2012; Tomasi & Volkow, 2012; Persson et al, 2014) as well as in age-related cognitive decline (Sheline et al, 2010; Buckner et al., 2005; Hafkemeijer et al, 2012). Other networks too have shown age-related changes: the auditory network shows age-related dedifferentiation, both in activity and in connectivity within itself and with other networks (such as DMN and salience network) (Hwang et al., 2007; Onoda et al, 2012). This age-related decline is observed in normal-hearing older adults, but is also associated with age-related hearing loss (Hwang et al., 2007; Onoda et al, 2012; Fitzhugh et al., 2019). The dopaminergic reward system shows decreased density with age in both striatal and extrastriatal regions (Erixon-Lindroth et al., 2005; Li & Rieckmann, 2014). Age-related decrease in dopaminergic function is linked to behavioral changes in a variety of cognitive and affective tasks. Cognitive changes include age-related declines with working memory capacity and maintenance (Landau et al., 2009; Klostermann et al., 2012), as well as with reward learning from prediction errors (Chowdhury et al., 2013; Samanez-Larkin et al., 2014). Reward learning is particularly of interest as it lies at the intersection of cognitive and affective tasks: the ability to form representations of predictions and prediction errors, which is fundamental to reward learning (Schultz et al., 1997), may be thought of as a cognitive process. On the other hand, the ability to represent valence and arousal of stimuli to be learned, and to form value-based representations of rewards more generally, may be considered more affect-based processing (Mather, 2016). While age-related decline has been observed for both cognitive and affective aspects of reward learning (Bäckman et al., 2010; Eppinger, Nystrom, & Cohen, 2012; Samanez-Larkin et al., 2014; Yee et al., 2019), some studies have also shown no age-related differences in activity of the reward network, such as during the processing of monetary incentives (Spaniole et al., 2015; Geddes et al., 2018), while other work has shown a more pronounced age-related decline in the memory-based, more cognitive aspects of reward learning, but a relative sparing of its more affective aspects (Samanez-Larkin et al., 2014; Geddes et al., 2018). At a behavioral level, older adults have also been reported to show more positive affect overall (Kessler & Staudinger, 2009), as well as less cognitive engagement during emotion-regulation tasks (Scheibe et al., 2015).

Given these findings, it would be of great interest to learn how age-related changes in functional network dynamics interact with reward responses to cognitively and affectively engaging sensory stimuli. In that regard, music has a unique power to elicit moments of intense emotional and physiological responses (Harrison & Loui, 2014), experiences that are intrinsically rewarding for most people (Zatorre, 2015). While music listening habits are known to be linked to psychological well-being in late adulthood/or late life (Laukka, 2006), the relationship between aging and the reward response to music is not well understood. On one hand, music-based interventions are hypothesized to be more enjoyable than other, non-musical cognitive training programs for older adults (Sutcliffe et al, 2020). Active music training in old age, such as through learning to play a musical instrument, could be a means to mitigate cognitive decline by facilitating auditory processes as well as more central processes such as cognitive reserve (Zendel & Alain, 2012; Alain et al, 2014; Ai et al, 2022). On the other hand, older adults show less sensitivity to reward (Eppinger, Nystrom, & Cohen, 2012), as subserved by differences in the reward system during gambling tasks (Geddes et al, 2018). Behavioral studies have also shown that sensitivity to musical reward, as assessed by self-report through the Barcelona Music Reward Questionnaire (BMRQ; Mas-Herrero et al., 2013), is lower in older adults than in younger adults (Belfi et al, 2021; Cardona et al, 2022). Addressing these gaps in our understanding of age-related differences in reward and functional network dynamics will have implications for the design of music-based interventions for healthy aging, as various strategies employed during music listening, even among receptive music-based interventions (e.g. listening for relaxation, imagery, or engagement), may cater to distinct brain networks that are differentially impacted by advancing age (Wheeler, 2015).

Cross-sectional functional neuroimaging studies may help shed light on the intrinsic and task-related differences in music reward sensitivity in younger and older adults. During music listening, regions involved with auditory processing, including but not limited to superior temporal gyrus (STG) and Heschl’s gyrus (HG), coactivate with multiple areas over and above their intrinsic activity and connectivity at rest, resulting in a richly connected task-based network for music listening that covaries with liking and familiarity (Koelsch et al, 2005; Loui et al, 2012; Ellis et al, 2013; Quinci et al, 2022).Coactivated areas include lateral prefrontal networks important for auditory-motor and linguistic processes, as well as emotion and reward associated brain structures, including ventral portions of the medial prefrontal cortex (mPFC), orbitofrontal cortex, and the dorsal and ventral striatum (Ferreri et al, 2019; Gold et al, 2019; Salimpoor et al, 2011; 2013). The STG receives input from primary auditory regions (Yeterian & Pandya, 1998; Kaas & Hackett, 2000). It is structurally and functionally connected to the IFG and anterior insula (Loui et al, 2009; Frey et al, 2008; Wang et al, 2020), the latter of which also plays a role in reward system processing (Sachs et al, 2016; Ai et al, 2022). The STG also shows structural and functional connectivity to the nucleus accumbens (Loui et al, 2017; Martinez-Molina et al, 2016). The latter connects to regions including the mPFC, caudate, and substantia nigra, together constituting the reward system (Karlsgodt et al, 2015). Auditory connections to the reward system are hypothesized to encode hedonic responses to music (Zatorre, 2015; Belfi & Loui, 2020).

Despite our understanding of the reward system, much is still not known about the driving factor behind state- and trait-level differences in musical reward processing. Neuroimaging studies have found that individual differences in reward sensitivity to music listening are related to differences in white matter connectivity to and engagement of areas in the dopaminergic systems, specifically in the striatum (Sachs et al, 2016; Loui et al, 2017; Martinez-Molina et al, 2019; Salimpoor et al., 2011; Zatorre, 2015). While the case of musical anhedonia represents an extreme lack of sensitivity to musical reward, music listening preferences also vary in the general population by the listener and for different situations (e.g. Rentfrow et al, 2011; North & Hargreaves, 2004). Even within the same individual, differences in musical reward processing can arise based on the listener’s past experience with the piece, autobiographical memories related to the piece, and one’s overall appreciation for the intentionality of the performer or composer (Pereira et al., 2011, Alluri et al, 2017; Aydogan et al, 2018; Janata, 2009). Agency in music listening also impacts the processing of musical information, with self-selection of musical stimuli leading to more active engagement for listeners (Quinci et al, 2022), possibly due to predictive processes that can be more strongly engaged when listening to self-selected music (Vuust et al, 2022).

Here, we compare the functional connectivity of auditory and reward networks in a sample of older adults (OA; N = 24; aged 54-89) and younger adults (YA; N = 24; aged 18-23). We do this using a combination of resting state fMRI and fMRI collected during a music listening task. Our hypotheses are threefold. First, we expect well-liked music to elicit greater activity from auditory processing and reward associated brain regions. We expect this to be true for both young and older adults, and for differences in activity to arise between the two age groups. Second, we expect music listening to increase functional connectivity within and between auditory processing and reward associated brain regions as compared to rest. We expect this to be true of both younger adults and older adults, and that these connections will be particularly strong while listening to well-liked musical stimuli. Third and finally, we expect that connectivity within and between auditory and reward associated regions will vary with age. While we expect this effect to be present during both rest and music listening, we expect that age will modulate the connectivity of auditory and reward-associated regions in response to music, resulting in an age x task interaction effect.

We test these hypotheses through a combination of whole-brain linear regression, seed-based connectivity, and region-of-interest-based (ROI-to-ROI) connectivity approaches. Whole-brain linear regression is used to test our first hypothesis, providing a metric for general metabolic activity during music listening. Seed-based and ROI-to-ROI connectivity measures are used to test our second and third hypotheses. While seed-based connectivity is intended as a more global method for determining functional connectivity, determining how auditory and reward networks (taken as a whole) may connect to the rest of the brain, ROI-to-ROI connectivity is intended as a more hypothesis-driven approach to determining the connectivity among and between known hubs of the auditory and reward networks.

## Materials and Methods

### Participants

We recruited equal samples of older and younger adults guided by median sample sizes of neuroimaging studies published in top neuroimaging journals in 2017 (23 participants) and 2018 (24 participants) (Szucs and Ioannidis, 2020). Twenty-four adults between the ages of 18 and 23 (M=18.58, SD=1.21) were recruited as the young adult group through the Northeastern University student population (20 identified as female, 3 as male, 1 as non-binary) . They had to meet the following inclusion criteria: 1) were 18 years of age or older, 2) had normal hearing, and 3) and pass an MRI screening.

Twenty-eight older adults were recruited as part of a longitudinal study (Quinci et al., 2022). Recruitment for the older adults took place through community outreach as well as online recruitment engines such as including craigslist.org and BuildClinical.com. Participants were prescreened by telephone prior to participation and were included if they 1) were over 50 years old, 2) had normal hearing, 3) passed the Telephone Interview for Cognitive Status (TICS) with a score of ≥ 31/41 indicating no cognitive impairment, and 4) passed MRI screening.

Participants were excluded if they 1) changed medications within 6 weeks of screening; 2) had a history of psychotic or schizophrenic episodes; 3) had a history of chemotherapy within the past 10 years; or 4) experienced serious physical trauma or were diagnosed with a serious chronic health condition requiring medical treatment and monitoring within 3 months of screening. Of the screened participants, four were unable to complete the MRI portion of the study, resulting in a final sample of twenty-four older adults (13 identified as male, 11 as female) between the ages of 54 and 89 (*M*=66.67, *SD*=7.79).

Participants were compensated for their time through either payment or course credit. This study was approved by the Northeastern University Institutional Review Board. This study is preregistered at https://osf.io/zxd42.

### Procedure

All participants completed a pre-screening phone call before being enrolled to ensure they met the inclusion/exclusion criteria described above. All participants were also asked to provide the researcher with the names of six of their favorite songs.

The young adult group was recruited from Northeastern University for a single-session study in return for course credit. The older adult group was recruited as part of a longitudinal music-based intervention study that included an MRI, blood draw, battery of neuropsychological tests, and series of surveys (as described separately in Quinci et al., 2022). For the present analyses, we used fMRI data from the older adults that was collected before the music-based intervention took place. All participants filled out an online version of the Barcelona Music Reward Questionnaire (BMRQ) (Mas-Herrero et al, 2013) to assess individual differences in reward sensitivity. The BMRQ is a twenty-item questionnaire in which participants rate on a five-point scale the extent to which they agree with statements indicating their degree of musical reward across five dimensions: musical seeking, emotion evocation, mood regulation, sensory-motor, and social reward.

### Special Considerations due to COVID-19

For the younger adults, data collection occurred exclusively during the COVID-19 pandemic, between October 2021 and March 2022. Data collection for older adults began in July 2019 and finished in November 2021, with nine participants completing their visits during the pandemic.

For the collection of human subject data during the COVID-19 pandemic, data collection followed a Resumption of Research Plan (RRP) that was developed and approved in consultation with the Institutional Review Board (IRB) of Northeastern University (for more details on special considerations due to COVID-19, see Quinci et al, 2022).

### fMRI Task

The fMRI task consisted of 24 trials altogether. In each trial, participants were first presented with a musical stimulus (lasting 20 seconds), then they were given the task of rating how familiar they found the music to be (familiarity rating lasted 2 seconds), and how much they liked the music (liking rating also lasted 2 seconds). Musical stimuli for the MRI task consisted of 24 different audio excerpts, each of which was 20 seconds in duration. Each audio stimulus was from one of the following three categories: participant self-selected music (6/24 stimuli), researcher-selected music including well-known excerpts spanning multiple musical genres (10/24 stimuli) and novel music spanning multiple genres (8/24 stimuli). Stimuli were presented in a randomized order, and participants made ratings of familiarity and liking on the scales of 1 to 4: for familiarity: 1=very unfamiliar, 2=unfamiliar, 3=familiar, 4=very familiar; for liking: 1=hate, 2=neutral, 3=like, 4=love. Participants made these ratings by pressing a corresponding button on a button-box (Cambridge Research Systems) inside the scanner. Participants wore MR-compatible over-the-ear headphones (Cambridge Research Systems) over musician-grade silicone ear plugs during MRI data acquisition. The spatial mapping between buttons and the numerical categories of ratings were counterbalanced between subjects to reduce any systematic association between ratings and the motor activity resulting from making responses.

### Behavioral Data Analysis

The proportion of each liking rating (hate, neutral, like, love) was averaged across participants for each group (younger adults, older adults). A chi-square test was run on the 2 x 4 matrix relating the frequency of each liking rating (hate, neutral, like, love) for each group (younger adults, older adults). Familiarity ratings are not separately analyzed for this manuscript, as the relationship between liking and familiarity is investigated in a separate report in the lab (Kathios et al, 2023) and the general similarity between liking and familiarity on brain activity during music listening has been demonstrated (Pereira et al., 2011). The effects of both ratings in the auditory and reward systems have also been shown in a separate report in our lab (Quinci et al, 2022).

### fMRI Data Acquisition

Images were acquired using a Siemens Magnetom 3T MR scanner with a 64-channel head coil at Northeastern University. For task fMRI data, continuous acquisition was used for 1440 volumes with a fast TR of 475 ms, for a total acquisition time of 11.4 minutes. Forty-eight axial slices (slice thickness = 3 mm, anterior to posterior, z volume = 14.4 mm) were acquired as echo-planar imaging (EPI) functional volumes covering the whole brain (TR = 475 ms, TE = 30 ms, flip angle = 60°, FOV = 240mm, voxel size = 3 x 3 x 3 mm^3^). The resting state scan followed the same parameters and included 947 continuous scans, for a total scan length of approximately 7 and a half minutes. T1 images were also acquired using a MPRAGE sequence, with one T1 image acquired every 2400 ms, for a total task time of approximately 7 minutes.

Sagittal slices (0.8 mm thick, anterior to posterior) were acquired covering the whole brain (TR = 2400 ms, TE = 2.55 ms, flip angle = 8°, FOV= 256, voxel size = 0.8 x 0.8 x 0.8 mm^3^) (as described in Quinci et al., 2022).

### fMRI Data Analysis

#### Pre-processing

Task and resting state fMRI data were preprocessed using the Statistical Parametric Mapping 12 (SPM12) software (Penny et al, 2011) with the CONN Toolbox (Whitfield-Gabrieli & Nieto-Castanon, 2012). Preprocessing steps included functional realignment and unwarp, functional centering, functional slice time correction, functional outlier detection using the artifact detection tool, functional direct segmentation and normalization to MNI template, structural centering, structural segmentation and normalization to MNI template, and functional smoothing to an 8mm gaussian kernel (Friston et al., 1995). Denoising steps for seed-based connectivity analyses included white matter and cerebrospinal fluid confound correction (Behzadi et al, 2007), motion correction, global signal regression, and bandpass filtering to 0.008-0.09 Hz.

#### Univariate Whole-Brain Analyses

Behavioral responses from the fMRI task were imported into R Studio, and onset values were extracted for each trial based on the liking rating participants awarded each stimulus (hate, neutral, like, love). For each participant, data were converted from 4D to 3D images, resulting in 1440 scans. The model was specified using the following criteria: interscan interval = 0.475 seconds, microtime resolution = 16, microtime onset = 8, duration = 42 as to only include data from when the participant was listening to music. First-level main effects of each liking rating were obtained using the SPM12 toolbox. Second-level main effects were analyzed for each group (young and older adults) for each liking rating (hate, neutral, like, love) using a one-sample t-test. Between-group contrasts (young vs. older adults) were also run for each liking rating. Results were corrected at the voxel and cluster threshold of *p* < 0.05 (FDR-corrected). Second-level results were visualized using the CONN Toolbox (Whitfield-Gabrieli, & Nieto-Castanon, 2012).

#### ROI-ROI Analyses

ROI-ROI analyses were performed using 18 auditory network ROIs and 18 reward network ROIs derived from the CONN default atlas. The auditory network seed consisted of all subdivisions of bilateral superior temporal gyrus (STG), middle temporal gyrus (MTG), inferior temporal gyrus (ITG), and Heschl’s gyrus, and the reward network seed consisted of anterior cingulate, posterior cingulate, and bilateral insular cortex, frontal orbital cortex, caudate, putamen, pallidum, hippocampus, amygdala, and nucleus accumbens, as defined by Wang et al., 2020. Preprocessed ROI timeseries data were extracted from participants’ first-level SPM.mat files using the Marsbar Toolbox (Brett et al., 2002). Values associated with each liking rating (hate, neutral, like, love) were separated for each participant and averaged across all stimuli with that particular rating. For each liking rating, ROI time-series data were correlated across all auditory regions (auditory-auditory), across all reward regions (reward-reward), and between auditory and reward regions (auditory-reward).

Follow-up analyses of ROI-ROI connectivity included a repeated-measures MANCOVA for participants who used all four liking ratings during the music listening task (N = 20 OA, 18 YA), comparing groups in auditory-auditory, auditory-reward, reward-reward, auditory-mPFC, auditory-PCC, reward-mPFC, and reward-PCC connectivity across our four levels of musical liking. Covariates included gender, self-reported musical reward, and within-group age.

#### Seed-Based Connectivity Analyses

Seed-based connectivity analysis was performed using the CONN Toolbox (Whitfield-Gabrieli, & Nieto-Castanon, 2012). Above defined auditory and reward network ROIs were combined, resulting in one seed for each network. For both seeds, contrasts were drawn between task (Task>Rest), age groups (YA>OA), and interactions between age and task.

Further contrasts determined linear and quadratic effects of music liking (as determined by in-scanner ratings), and selection. Finally, to rule out any confounding effects of within-group age differences, gender, and musical reward as measured by the BMRQ, direct subject effect contrasts of these metrics were performed for each condition and seed effect. Main effect contrasts of group, task, selection, and liking were also repeated using within-group age and BMRQ as covariates of no interest, and relevant differences to the main contrasts are presented when applicable.

## Results

### Behavioral Results: Barcelona Music Reward Questionnaire

The Barcelona Music Reward Questionnaire (BMRQ) was higher in younger adults (M = 83.2, SD = 7.28) than in older adults (M = 72.3, SD = 9.97), as confirmed by an independent samples t-test (*t*(46) = 4.34, *p* < 0.001), indicating that younger adults experience greater self-reported music reward, replicating previous reports (Mas-Herrero et al, 2013; Belfi et al, 2020; Cardona et al, 2022).

### Frequency of Liking Ratings per Group

While both groups used all liking ratings, older adults used more extreme ratings: a Chi-squared test showed a significant association between group (younger adults vs older adults) and proportion of liking ratings for *χ*^2^(3) = 12.7, *p* = 0.005. Table 1 shows the proportion of each liking rating given by young and older adults respectively.

**Table 1.** Liking Rating Frequencies by Group

|  | Rating |  |  |  |
| --- | --- | --- | --- | --- |
|  | Hate | Neutral | Like | Love |
| **Young Adults** |  |  |  |  |
| % within Group | 13.80% | 32.50% | 23.20% | 30.50% |
| % within Rating | 43.60% | 57.90% | 46.30% | 48.90% |
| % of Total | 6.90% | 16.20% | 11.60% | 15.30% |
| **Older Adults** |  |  |  |  |
| % within Group | 17.80% | 23.50% | 26.80% | 31.90% |
| % within Rating | 56.40% | 42.10% | 53.70% | 51.10% |
| % of Total | 8.90% | 11.80% | 13.40% | 16.00% |
| **Total** |  |  |  |  |
| % of Total | 15.80% | 28.00% | 25.00% | 31.20% |

### Univariate Whole-Brain Results: Loved Music Activates Auditory and Reward Networks

For both young and older adults, music rated as hated, neutral, liked, and loved all activated auditory regions including the Heschl’s gyrus, planum temporale, planum polare, STS, STG, and medial temporal gyrus (MTG), as well as some regions outside the auditory network, specifically the hippocampus. For younger adults, music rated as hated and liked also activated the parahippocampal gyrus, and music rated as liked additionally activated the brainstem. When listening to loved music, several additional regions became active in both groups: the medial prefrontal cortex (mPFC), paracingulate gyrus, posterior cingulate cortex, precuneus, orbitofrontal cortex and lateral occipital cortex. Also during loved music, younger adults showed activity in the superior frontal gyrus, parahippocampal gyrus, brainstem, and supplementary motor area (SMA), whereas older adults showed activity in the ventral striatum/ nucleus accumbens, brainstem, and cerebellum (see Table 2 and Figure 1).

**Figure 1:**
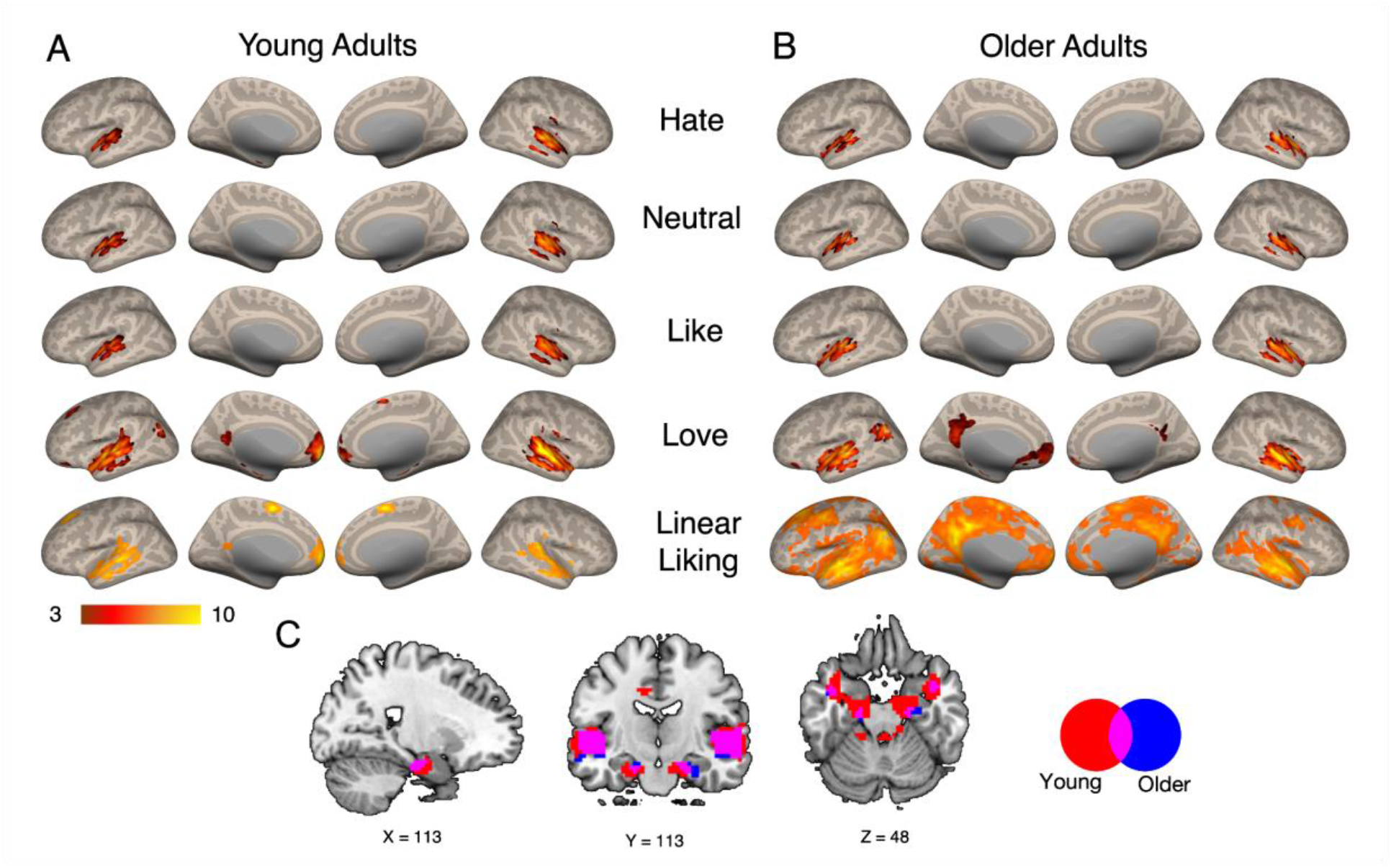
Functional Activity Across Liking Ratings. Univariate 2nd-level results showing activity at each level of liking (rows 1-4) and linear contrast of liking (row 5) for younger adults (A) and older adults (B) across liking ratings (p < 0.05 p-FDR corrected for voxel height and cluster size). (C) Subcortical structures involved with listening to loved music for younger adults (red), older adults(blue), and both (purple) (voxel height p < 0.05 p-FDR corrected; cluster size k > 10).

**Table 2.**
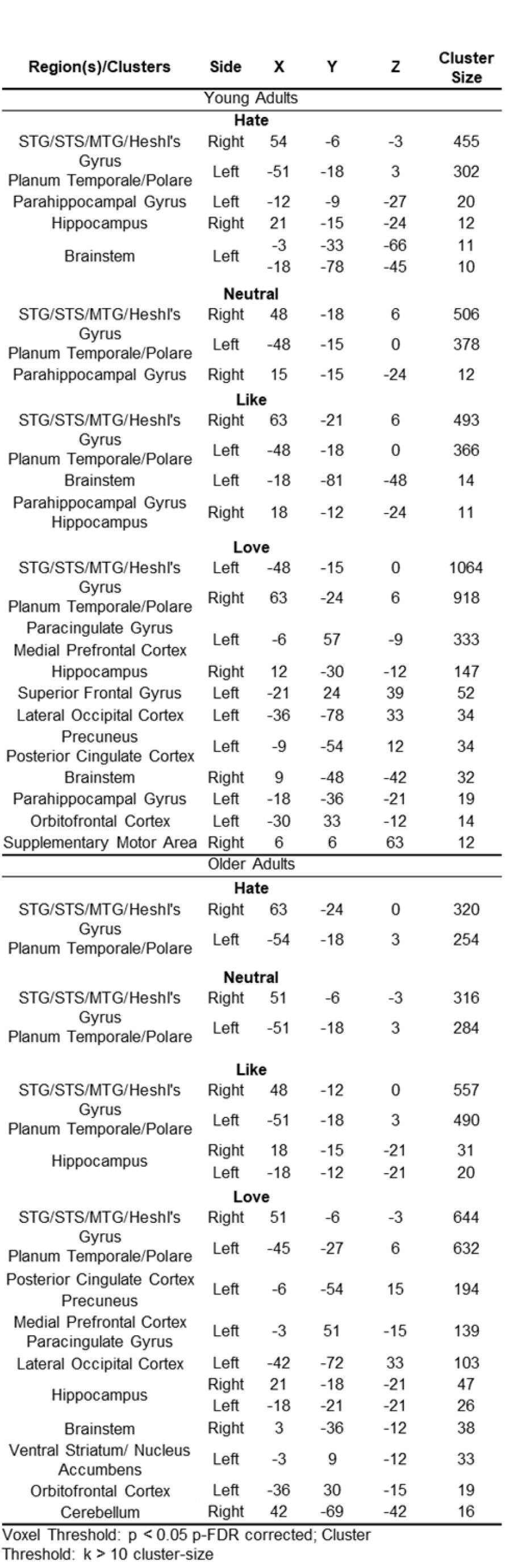
Univariate Whole-Brain Results

Young and older adults both showed a significant linear effect of liking rating (Hate, Neutral, Like, Love [-3 -1 1 3]) on activity across a broad range of regions, including auditory cortex, mPFC, PCC, and striatum (Figure 1 A-B, bottom row). While linear-liking effects were present in both groups, OA showed larger clusters of significance spanning a greater portion of the brain, while YA showed more regional specificity of the effect. Follow-up testing indicated no significant quadratic effect of liking (Hate, Neutral, Like, Love [1 -1 -1 1]) for either group, and whole-brain comparisons for young vs. older adult contrasts showed no significant effects of group at the p < .05 FDR-corrected level.

### ROI-ROI Connectivity: Linear and Quadratic Relationships with Liking and Effects of Age Group

Results of our ROI-ROI analysis are summarized in Figure 2. Three repeated-measures ANOVAs were run to evaluate the main effects of liking ratings and age and the rating x age interaction for connectivity among auditory and reward regions, and between auditory and reward regions. We also ran linear and quadratic contrasts to further investigate the relationship between ratings and connectivity for the conditions listed above.

**Figure 2:**
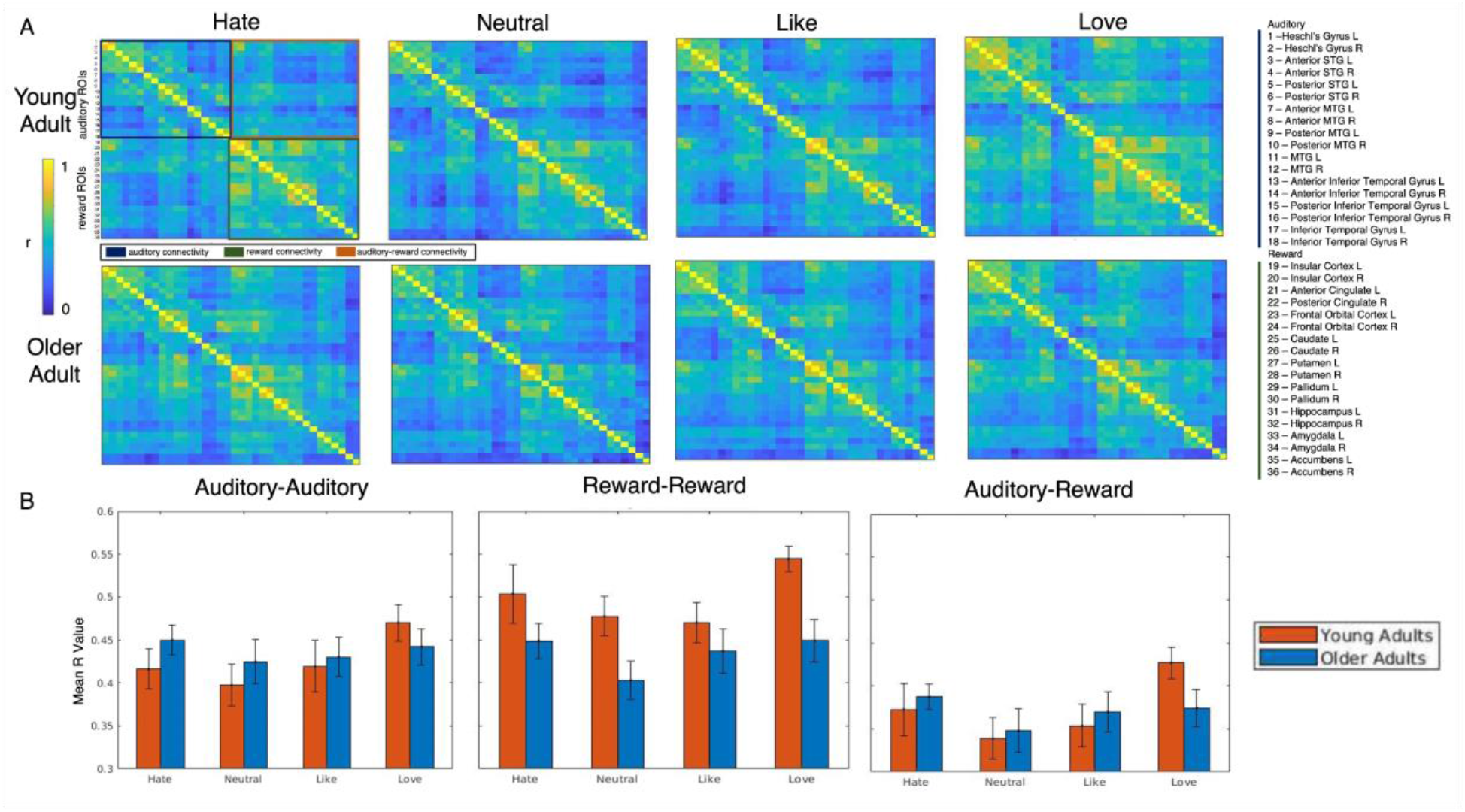
Age Related Differences in ROI-ROI Connectivity for Various Levels of Liking. (A) Correlation matrices showing the relationship between auditory and reward regions for different liking ratings for young and older adults. (B) Bar graphs depicting overall connectivity between auditory-auditory, reward-reward, and auditory-reward regions for young (orange) and older (blue) adults for hated, neutral, liked, and loved musical stimuli.

For connectivity among auditory regions, there was a significant main effect of liking (F(3, 108) = 3.06, p = 0.032, *η_p_*^2^= 0.078). Bonferroni corrected pairwise comparisons revealed that beta-weights for stimuli rated as loved were significantly greater than those rated as neutral. There was no significant main effect of age group (F(1, 36) = 0.804, p = 0.376, *η_p_*^2^ = 0.022) and no significant rating x age group interaction (F(3, 108) = 0.807, p = 0.492, *η_p_*^2^ = 0.022). While the linear contrast was only marginally significant (F(1, 36) = 3.42, p = 0.073, *η_p_*^2^ = 0.087), there was a significant quadratic contrast (F(1, 36) = 6.95, p = 0.012, *η_p_*^2^ = 0.162), suggesting that hated and loved music had greater functional connectivity compared to neutral and liked music.

For connectivity within reward regions, there was a significant main effect of liking (F(3,108) = 3.19, p = 0.027, *η_p_*^2^ = 0.081). Bonferroni-corrected pairwise comparisons revealed that beta-weights for stimuli rated as loved were significantly greater than those rated as neutral and those rated as liked. There was a significant main effect of age (F(1,36) = 5.13, p = 0.030, *η_p_*^2^ = 0.125), suggesting that younger adults have greater connectivity within reward regions compared to older adults. There was no significant rating x age interaction (F(1,108) = 1.43, p = 0.237, *η_p_*^2^ = 0.038). There was no significant linear contrast (F(1, 36) = 2.28, p = 0.139, *η_p_*^2^ = 0.060), but there was a significant quadratic contrast (F(1, 36) = 8.26, p = 0.007, *η_p_*^2^ = 0.187). Thus, hated and loved music had greater connectivity compared to neutral and liked music.

For connectivity between auditory and reward regions, there was a significant main effect of liking (F(3, 108) = 3.60, p = 0.016, *η*^2^ = 0.091). Bonferroni-corrected pairwise comparisons revealed that beta-weights for stimuli rated as loved were significantly greater than those rated as neutral. There was no significant main effect of age (F(3, 36) = 0.002, p = 0.962, *η_p_*^2^ <0.001) and no significant rating x age interaction (F(3, 108) = 1.49, p = 0.221, *η_p_*^2^ = 0.040).

There was both a significant linear contrast (F(1, 36) = 4.20, p = 0.048, *η_p_*^2^ = 0.104) and a significant quadratic contrast (F(1, 36) = 7.28, p = 0.011, *η_p_*^2^ = 0.168); in other words, auditory-reward connectivity scaled both with liking and with valence.

### Seed-Based Connectivity: Age Group Differences in Auditory and Reward Network Connectivity

We seeded the auditory and reward networks separately and investigated the effects of age, task, musical liking ratings, selection, and BMRQ on functional connectivity from these seed networks. All seed-based analyses presented were performed at the p < 0.05, FDR-corrected level for both voxel height and cluster size. Significant clusters across all analyses are summarized in Table 3. Statistical maps of all significant results can be found at https://neurovault.org/collections/QSMRNLRW/

**Table 3.**
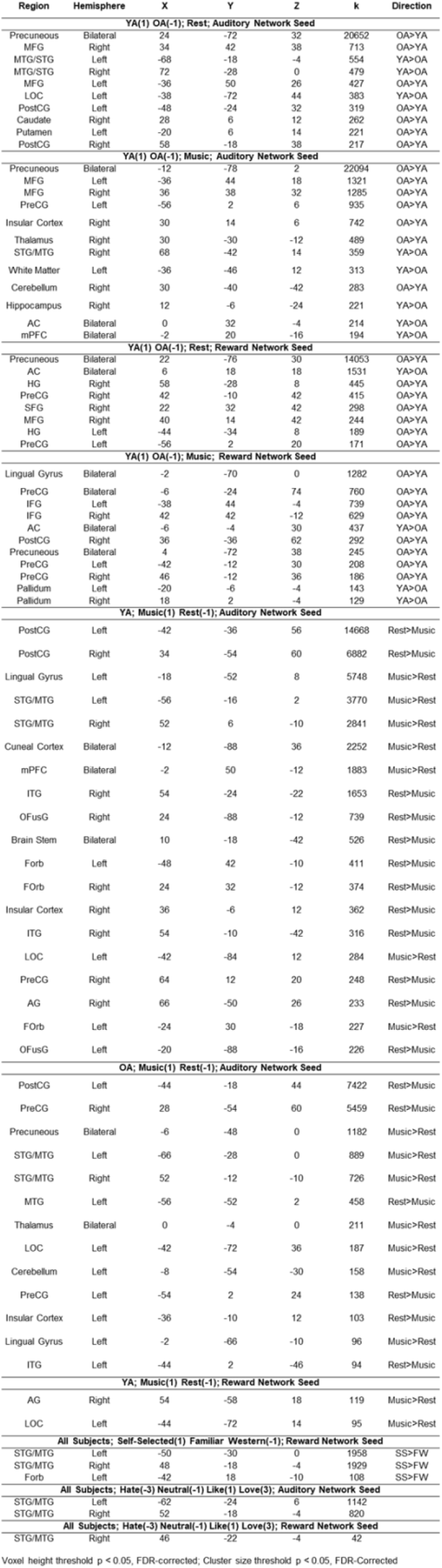
Seed Based Connectivity Results

### Lower Network Differentiation in Older Adults

Contrasts between older and younger adults (YA>OA) revealed significant effects of auditory and reward network-seeded connectivity during both music listening and rest. Younger adults showed significantly higher auditory network-seeded connectivity to other regions in the auditory network, including bilateral STG and MTG, during rest (Figure 3A-B). During music listening, these effects were limited to the right hemisphere, and younger adults additionally showed higher connectivity between auditory network and mPFC, hippocampus, and anterior cingulate compared to older adults (see Figure 3C-D). Older adults showed higher connectivity to somatosensory, reward, and posterior default-mode processing regions compared to younger adults, for which effects were similar during music listening as during rest. When including BMRQ as a covariate of no interest, significant effects were more constrained (see Supplementary Figure 1). During rest, regions of higher auditory network connectivity for the YA group were limited to the right hemisphere, and for the OA group, higher connectivity in the somatosensory and frontal clusters no longer reached significance. During music listening, only higher connectivity in the precuneus for the OA group remained significant, suggesting that the other effects are explained by individual differences in BMRQ.

**Figure 3:**
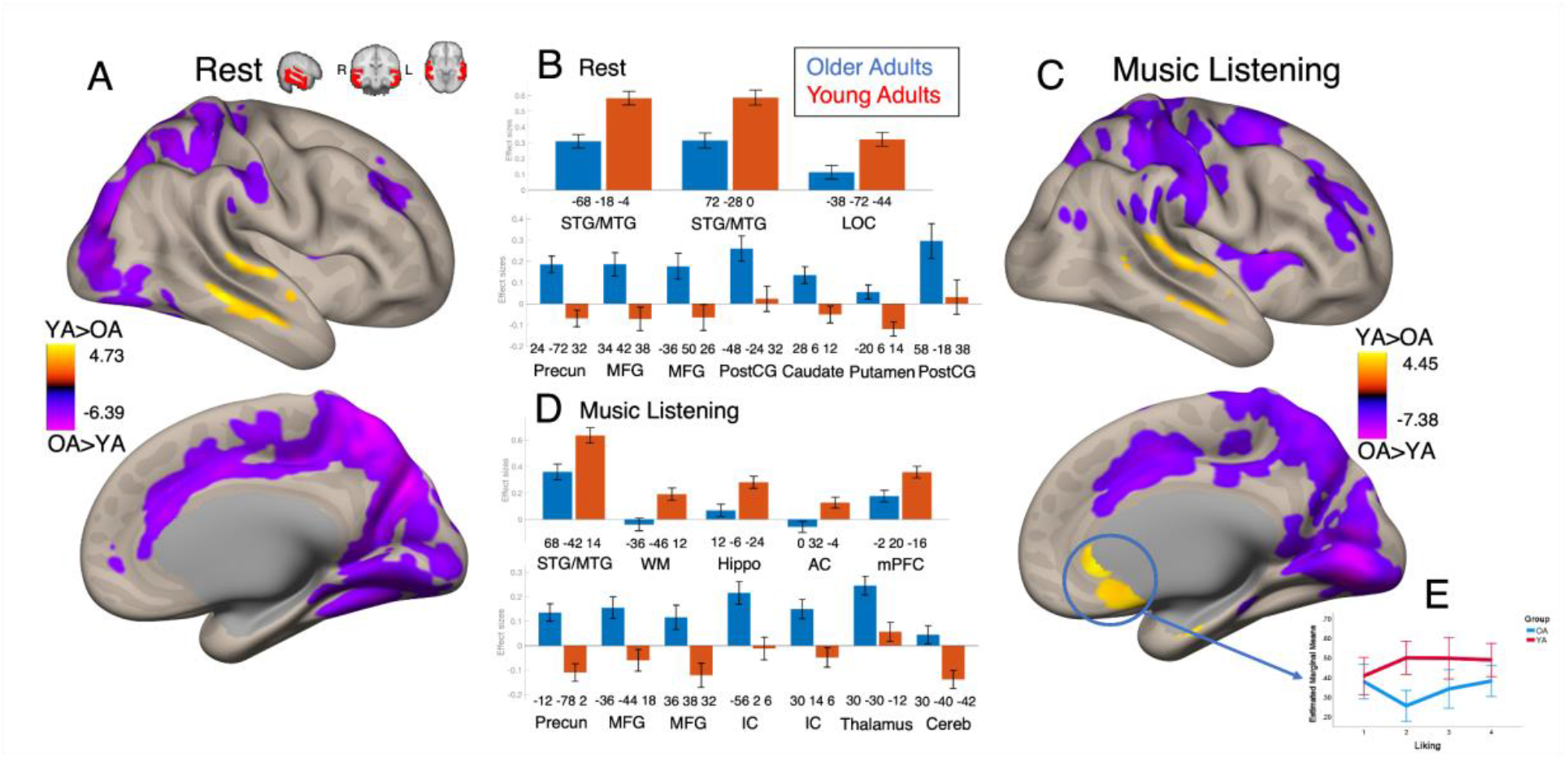
Age Differences in Auditory Network Seed-Based Connectivity: Significant clusters for YA>OA contrast in auditory network seed, significant at the p < 0.05, FDR-corrected level for both voxel height and cluster size. (A) Visualization of significant clusters in right hemisphere during resting state scan, including clusters favoring younger adults (warm tones) and clusters favoring older adults (cool tones). (B) Bar graphs depicting effect sizes for all significant clusters in resting state contrast. Clusters with higher connectivity in younger adults (red) are depicted in the top graph, and clusters with higher connectivity in older adults (blue) are depicted in the bottom graph. Cluster labels are MNI coordinates for center of mass of each significant cluster, as well as the most prevalent anatomical region included within each cluster. (C) Visualization of significant clusters in right hemisphere during music listening task, including clusters favoring younger adults (warm tones) and clusters favoring older adults (cool tones). (D) Bar graphs depicting effect sizes for all significant clusters in music listening contrast. Clusters with higher connectivity in younger adults (red) are depicted in the top graph, and clusters with higher connectivity in older adults (blue) are depicted in the bottom graph. (E) Line graph depicting ROI-ROI connectivity between auditory network regions and mPFC comparing across groups and liking ratings. Error bars depict +/- two standard errors, and marginal means are adjusted for covariates of gender, BMRQ, and within group age.

At rest, younger adults showed significantly higher reward-network seeded connectivity to anterior cingulate (AC) while older adults showed significantly higher connectivity to posterior DMN and somatosensory regions (Figure 4A-B). During music listening, younger adults once again showed connectivity to reward-associated regions, now including both AC and bilateral pallidum, while older adults now showed significant clusters of increased connectivity in bilateral IFG, in addition to effects observed at rest (Figure 4C-D).

**Figure 4:**
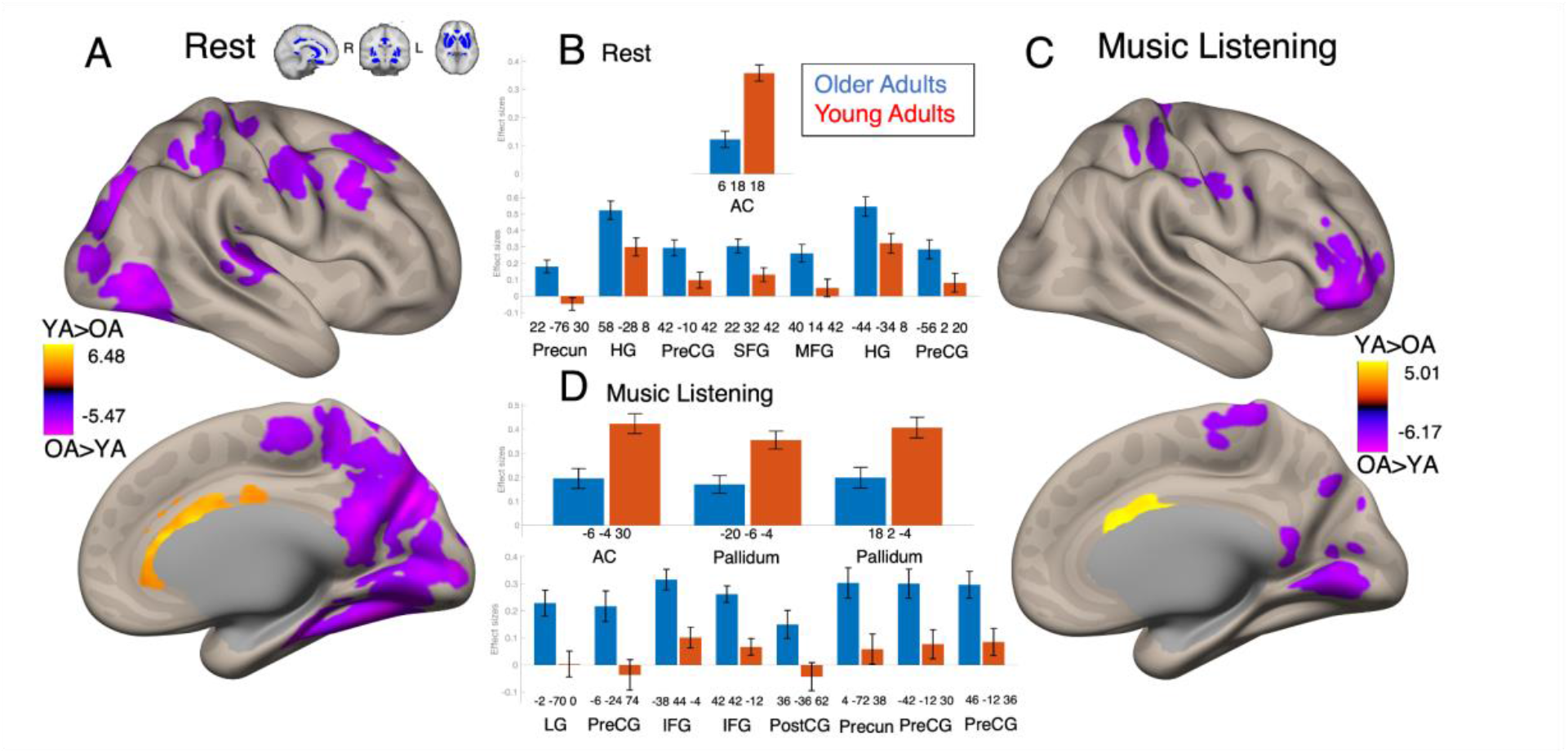
Age Differences in Reward Network Seed-Based Connectivity: Significant clusters for YA>OA contrast in reward network seed, significant at the p < 0.05, FDR-corrected level for both voxel height and cluster size. (A) Visualization of significant clusters in right hemisphere during resting state scan, including clusters favoring younger adults (warm tones) and clusters favoring older adults (cool tones). (B) Bar graphs depicting effect sizes for all significant clusters in resting state contrast. Clusters with higher connectivity in younger adults (red) are depicted in the top graph, and clusters with higher connectivity in older adults (blue) are depicted in the bottom graph. Cluster labels are MNI coordinates for center of mass of each significant cluster, as well as the most prevalent anatomical region included within each cluster. (C) Visualization of significant clusters in right hemisphere during music listening task, including clusters favoring younger adults (warm tones) and clusters favoring older adults (cool tones). (D) Bar graphs depicting effect sizes for all significant clusters in music listening contrast. Clusters with higher connectivity in younger adults (red) are depicted in the top graph, and clusters with higher connectivity in older adults (blue) are depicted in the bottom graph.

When including BMRQ as a covariate of no interest, only the AC cluster showing higher connectivity in the YA group at rest and the IFG cluster showing higher connectivity in the OA group during music listening remained significant (see Supplementary Figure 2).

Considering differences in age range across our two samples, all seed-based connectivity analyses were repeated using age, demeaned based on mean age for each sample, as a covariate of no interest. In all cases, there were no notable differences in the observed main effects of group or task. There were also no significant main effects of within-group age across all conditions.

### Higher Auditory-mPFC Connectivity in Younger Adults during Music Listening: Effects of Task

Auditory network-seeded connectivity showed significant differences between task and rest within both of our age groups. In younger adults, connectivity was higher during music listening than rest in auditory cortex, reward, and default-mode regions, and at rest, connectivity was most notably higher than music listening in somatosensory regions and portions of the ITG (Figure 5A-B). In older adults, many of the observed effects were similar, although positive effects of music listening were generally more limited, and there was no increased connectivity observed within the mPFC (see Figure 5C-D). Reward network-seeded main effects of task were only observed in the YA group, including an increase in connectivity to bilateral lateral occipital cortex (LOC) and angular gyrus (AG) during music listening (see Figure 5E). Inclusion of BMRQ as a covariate of no interest resulted in no significant clusters for this contrast. No age by task interactions were observed.

**Figure 5:**
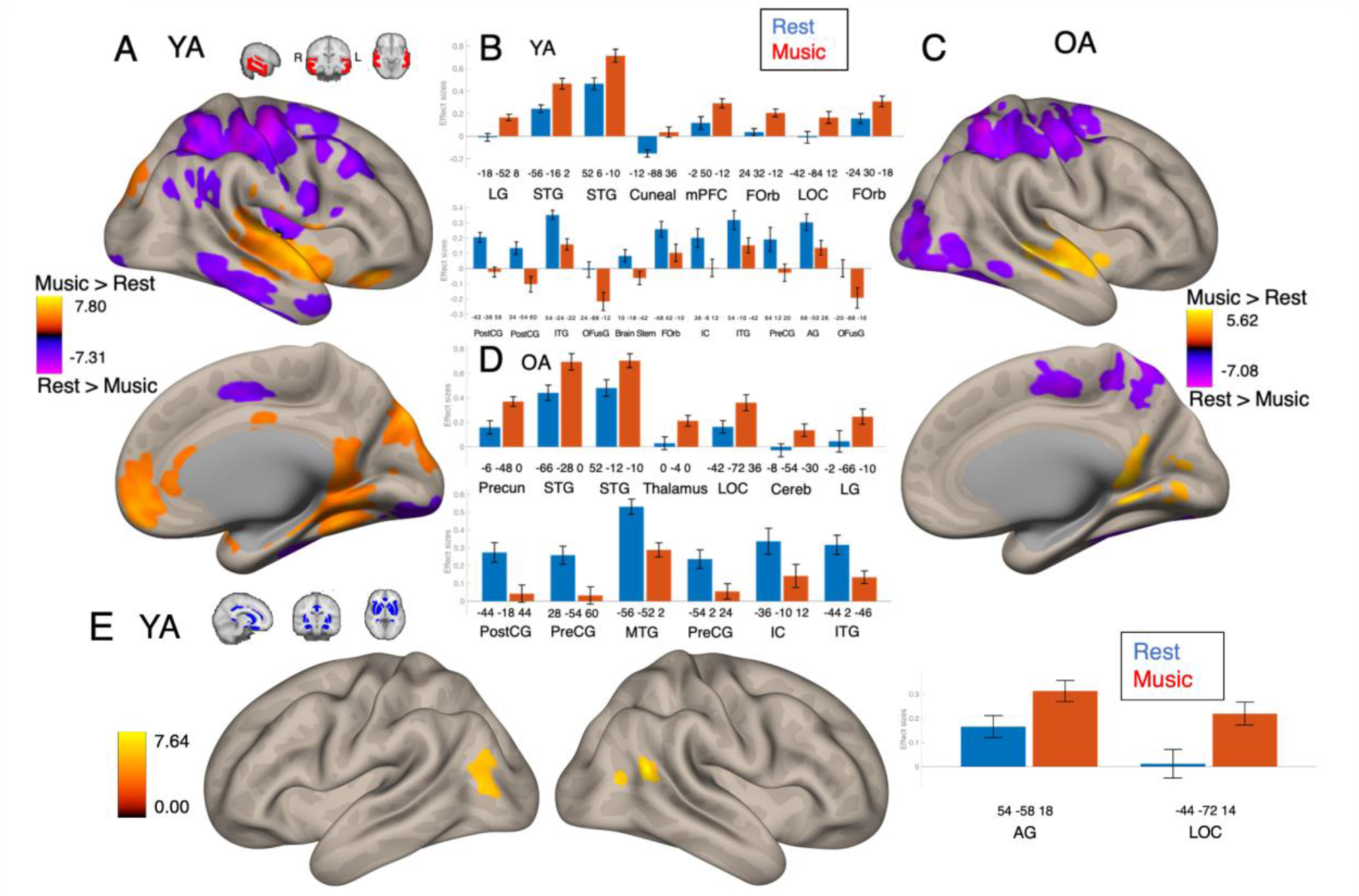
Task Differences in Seed-Based Connectivity: Significant clusters for music > rest contrast, significant at the p < 0.05, FDR-corrected level for both voxel height and cluster size. (A) Visualization of significant clusters from auditory network seed in right hemisphere of younger adults, including clusters favoring music listening (warm tones) and clusters favoring rest (cool tones). (B) Bar graphs depicting effect sizes for all significant clusters in young adult contrast. Clusters with higher connectivity during music listening (red) are depicted in the top graph, and clusters with higher connectivity during rest (blue) are depicted in the bottom graph. Cluster labels are MNI coordinates for center of mass of each significant cluster, as well as the most prevalent anatomical region included within each cluster. (C) Visualization of significant clusters from auditory network seed in right hemisphere of older adults, including clusters favoring music listening (warm tones) and clusters favoring rest (cool tones). (D) Bar graphs depicting effect sizes for all significant clusters in older adult contrast. Clusters with higher connectivity during music listening (red) are depicted in the top graph, and clusters with higher connectivity during rest (blue) are depicted in the bottom graph. (E) Visualization of significant clusters from Reward network seed in younger adults, and bar graphs depicting effect sizes for all significant clusters. Cluster labels are MNI coordinates for center of mass of each significant cluster, as well as the most prevalent anatomical region included within each cluster.

### Well-liked and Self-Selected Music Increases Auditory-Reward Connectivity: Musical Stimulus Effects

Effects contrasting different musical stimuli were also observed. There was significantly higher connectivity across all subjects between reward network and bilateral auditory cortex in the participant-selected music condition as compared to researcher-selected familiar western music (see Figure 6A). There was also a significant linear effect of musical liking ratings across all subjects in auditory network seed. This took the form of increased connectivity to auditory regions, including bilateral STG and MTG (see Figure 6B). Inclusion of BMRQ as a covariate of no interest resulted in additional significant clusters for this effect in the amygdala and brainstem (Supplementary Figure 3). Reward network-seeded connectivity to auditory cortex also showed a significant linear effect of liking; however this effect was limited to right lateral STG (see Figure 6C). There were no significant quadratic effects of liking observed, and no significant stimulus x group interaction effects.

**Figure 6:**
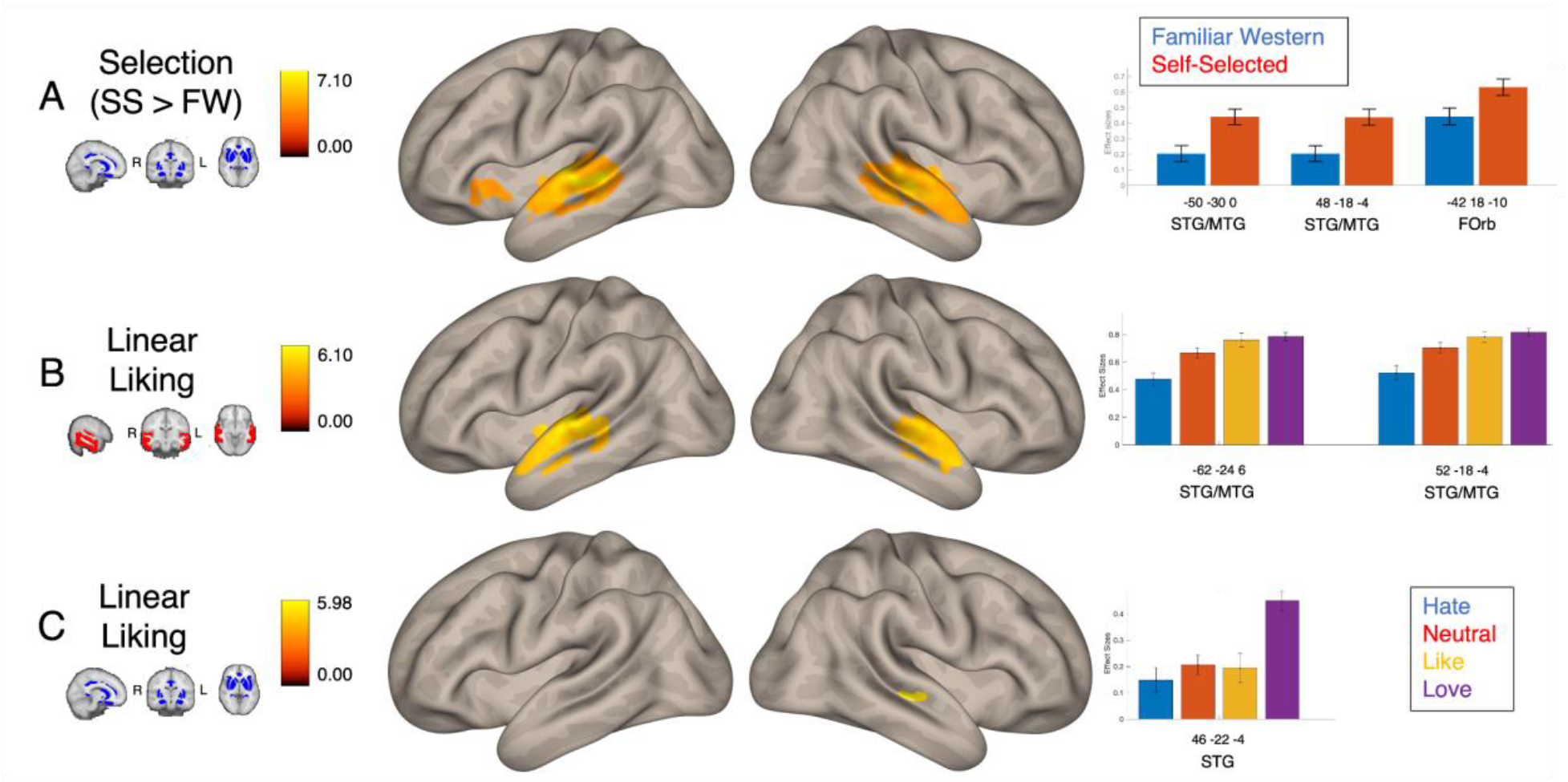
Differences of Musical Preference in Seed-Based Connectivity: Significant clusters for effects of musical selection (self-selected > familiar western) and linear effects of musical liking in auditory and reward network seed-based connectivity ([-3,-1,1,3], two-tailed T-test; p < 0.05, FDR-corrected for voxel height and cluster size). (A) Visualization of significant clusters from reward network seed showing effects of selection, and bar graphs depicting effect sizes. Cluster labels are MNI coordinates for center of mass of each significant cluster, as well as the most prevalent anatomical region included within each cluster. (B) Visualization of significant clusters from auditory network seed showing linear effects of liking, and bar graphs depicting effect sizes. Cluster labels are MNI coordinates for center of mass of each significant cluster, as well as the most prevalent anatomical region included within each cluster. (C) Visualization of significant clusters from reward network seed showing linear effects of liking, and bar graphs depicting effect sizes. Cluster labels are MNI coordinates for center of mass of each significant cluster, as well as the most prevalent anatomical region included within each cluster.

There were no significant main effects of BMRQ or gender for either seed network in rest and music listening, and no significant BMRQ x task or gender x task interaction effects.

### Linear and Quadratic Effects of Auditory, Reward, and DMN Network Connectivity for Liking and Age

*Follow-up MANCOVA of connectivity among and between auditory network ROIs, reward network ROIs, mPFC, and PCC (see Supplementary Figure 4) indicated a significant main effect of group (F(1, 36) = 4.21, p = 0.003, η_p_*^2^ *= 0.522), characterized by higher connectivity in YA relative to OA. Also observed were quadratic, but not linear, interactions between liking and group in auditory-mPFC (F(1, 36 = 4.31, p = 0.046, η_p_*^2^ *= 0.116; Figure 3E) and auditory-PCC connectivity (F(1, 36) = 5.57, p = 0.024, η_p_*^2^ *= 0.144), as well between liking and BMRQ in auditory-auditory (F(1, 36) = 6.03, p = 0.010, η_p_*^2^ *= 0.155), auditory-mPFC (F(1, 36) = 7.357, p = 0.011, η_p_*^2^ *= 0.182), and auditory-PCC connectivity (F(1, 36) = 4.27, p = 0.047, η_p_*^2^ *= 0.115). Linear interactions between liking and within-group age were also observed in auditory-auditory (F(1, 36) = 5.02, p = 0.032, η_p_*^2^ *= 0.132) and auditory-reward connectivity (F(1, 36) = 4.53, p = 0.041, η_p_*^2^ *= 0.121)*.

## Discussion

Here we characterize for the first time age-related differences in the neural substrates of musical reward. These data provide insight into the mechanisms underlying observed differences in musical reward in young and older adults (Belfi et al., 2020; Cardona et al, 2022; Mas-Herrero et al, 2013), relate the contributions of auditory and reward networks to music listening at varying levels of musical preference, and provide potential brain-based targets for the development of music-based interventions in older adult populations.

Univariate analyses showed that both young and older adults activated multiple regions within the auditory network during music listening, as well as the mPFC during music listening that was rated as loved. The engagement of mPFC when listening to preferred music is consistent with its observed role in musical reward processing (Salimpoor et al, 2013). Both groups showed specific activation of the hippocampus during preferred music listening.

Although not classically considered part of the mesolimbic reward pathway, the hippocampus is an input into the ventral striatum (Lebreton et al, 2009) and is known to respond to reward and value, especially during reward learning (Davidow et al, 2016; Wimmer et al, 2018), memory formation, and emotion processing (Bannerman et al, 2004; Frühholz et al, 2014). Here, the majority of main effects of music listening were similar between groups, suggesting that the overall activity patterns associated with music listening is preserved during aging. Nevertheless, univariate results, especially when listening to loved music, showed some clusters that were only significant in one group or another. Although a between-groups contrast was not statistically significant, main effects of music listening in each group pointed to differences when listening to loved music. Older adults showed significant precuneus activity whereas younger adults showed significant SMA activity. The precuneus is a highly metabolically active area in the brain; while being a hub of the DMN, it is also tied functionally to episodic memory retrieval as well as the experience of agency (Cavanna & Trimble, 2006; Darby, Burke & Fox, 2018). In this study, the finding of larger clusters of activity in the precuneus that scale with liking in older adults may reflect higher metabolic demands in older adulthood for music listening; alternately, it may reflect default mode functions including but not limited to episodic memory processing and/or a greater sense of agency during music listening.

Only younger adults showed SMA activity when listening to loved music. SMA is often activated in auditory imagery (Lima et al; 2016), especially during beat perception in rhythmic music (Nombela et al, 2013). Neurophysiological responses comparing young and older adults during music listening have shown that both groups are broadly sensitive to the beat in neural phase locking measures, but subtle differences persist in topographical distributions of phase locking at different levels of the beat (Tichko et al, 2022).

In ROI-ROI analyses, well-liked music resulted in higher connectivity among and between auditory and reward regions, as compared to music rated neutrally. The difference between young and older adults in functional connectivity was especially accentuated when listening to preferred music. We see that connectivity of auditory and reward regions during music listening is not a linear function in preference for the music, but rather determined by a u-shaped relationship with liking ratings. This stands in contrast to the effect of liking observed in our seed-based connectivity analysis, in which auditory network seed showed a significant linear effect, but not a quadratic effect, in connectivity to specific auditory regions. We interpret these different effects as resulting from differences in the methodologies of SBC and ROI-ROI analyses. SBC started with the auditory and reward networks taken as a whole, and thus we observed that the strongest connections arising within these networks lie within regions that show a linear effect of liking. Meanwhile, our ROI-ROI approach granted equal weight to each region identified as contributing to the auditory and reward network. In this regard, vertices showing strong linear effects of liking were accompanied by vertices that showed stronger connectivity during hated music, resulting in the observed U-shaped curve.

We also observed higher overall connectivity among reward regions in younger adults as compared to older adults. This is in keeping with observed decreases in overall hedonic response with age (Eppinger, 2012), and are consistent with previously observed losses of functional network connectivity in older adults, both generally (Sala-Llonch, Bartrés-Faz & Junqué, 2015; Sheline et al, 2010) and specific to reward sensitive systems (Geddes et al, 2018).

Seed-based connectivity results further support the finding that younger adults have stronger within-network seed-based connectivity than older adults. This was seen in auditory-auditory and reward-reward connections during both rest and music listening. However, music listening and individual differences in musical reward did mediate these effects. Auditory-auditory group effects were limited to the right hemisphere during the music listening task, as well as when BMRQ was included as a covariate of no interest during rest. Research into hemispheric differences in auditory processing has long supported a right-hemispheric dominance in music processing as compared to speech (Zatorre, 1992). Musical training can also lead to a reduction in the rightward asymmetry of musical feature processing (Ono et al, 2011). As such, one possible explanation is that primary musical processing regions in the right hemisphere show a more robust effect of age, while left hemispheric differences can be accounted for by differences in task and individual differences in musical reward.

Alternatively, left auditory cortex is known to be more sensitive to rapid temporal information, whereas the right auditory cortex is more sensitive to spectral information (Zatorre & Belin, 2001; Boemio et al, 2005; Albouy et al, 2020). These differences are also reflected in the way people listen to music: individuals who focus their listening towards spectral features show higher activity in right HG, whereas those who focus on more holistic features show more left-lateralization (Schneider et al, 2005). Therefore, another potential interpretation of these results, that may be tested in future studies, is that age-related differences affect spectral processing or spectral acuity more than holistic or temporal auditory processing, the latter being explained by differences in musical reward and recovered by active music listening.

Furthermore, inclusion of BMRQ during the music listening task removed all auditory-auditory and reward-reward effects favoring younger adults. These results suggest that individual differences in musical reward account for some of the observed differences between age groups, and that music listening may remediate age-related deficits in auditory and reward network coherence.

Turning now to specific connectivity effects between our seed networks and other, out-of-network regions, one major finding is that music listening led to greater connectivity of mPFC from auditory network in younger adults as compared to older adults. This can be observed in both the task > rest contrast, in which the effect was seen in younger but not older adults, and YA > OA contrast, in which the effect was seen during music listening, but not rest. This, taken together with findings from whole brain regression, suggests that while mPFC is activated during preferred music listening across age groups, the inclusion of mPFC in the auditory system during music listening in general is more readily observable in young adults specifically. This is also supported by the results of the follow-up MANCOVA comparing the average pairwise correlations between auditory regions and mPFC, which resulted in a significant group by liking interaction. This is consistent with previous work demonstrating that functional connectivity of the default-mode network, and especially mPFC, decreases with age (Sambataro et al, 2010; Staffaroni et al, 2018), and continues to decline with age-related illnesses such as Alzheimer’s disease (Hafkemeijer et al, 2015; Schouten et al, 2016; Staffaroni et al, 2018).

Another region that showed a significant interaction between music listening and age was the lateral occipital cortex (LOC). Auditory-seeded connectivity to LOC was higher in music listening than rest in both groups; however only the YA group showed increased connectivity between reward network and LOC. This result was not expected, considering LOC’s primarily observed role in object perception (Malach et al, 2002; Nagy et al, 2012). However, some work has shown preferential connectivity between precuneus and LOC in trained musicians (Tanaka & Kirino, 2016) and increased activity and structural connectivity of LOC during music reading as compared to word reading (Bouhali et al, 2020), suggesting a potential role of LOC in music processing that is not yet fully understood.

Meanwhile, older adults showed greater connectivity between auditory-seeded and reward-seeded networks and out-of-network regions, including MFG, precentral gyrus, and postcentral gyrus, consistent with the notion of age-related dedifferentiation. Older adults also showed significantly higher connectivity to precuneus, lending further support to the effect observed within the global linear regression model. For the most part, effects favoring older adults took the form of moderately positive effect sizes in the OA group compared to moderately negative or near-zero effect sizes in the YA group. This suggests that while older adults show a decrease of network coherence, there is simultaneously a gain of more distributed connectivity between functionally distinct brain regions.

However, some of the specific effects favoring older adults do warrant further discussion. For one, older adults showed higher connectivity between reward network and bilateral IFG as compared to younger adults that was specific to the music listening task, and the right-lateral component of this effect was the only group effect within that contrast to survive correction for differences in BMRQ. IFG is known to play a role in musical syntax processing (Koelsch et al, 2005; Kunert et al, 2015), and right IFG in particular is dominant over left IFG for the perception of pitch in song (Merrill et al, 2012) and for the integration of harmonic information into auditory and motor systems (Bianco et al, 2016). The fact that this result is only observed during music listening may suggest that the IFG is specifically responding to musical stimuli, and the effect favoring older adults may reflect compensatory changes in aging, and suggest that older adults may be more efficiently using IFG to incorporate musical information into multiple systems, including the reward network.

Another region that showed increased connectivity to auditory and reward networks in older adults is the lingual gyrus (LG). LG plays a critical role in the encoding and retrieval of visual memories (Bogousslavsky et al, 1987; Machielsen et al, 2000). It has also shown increased structural and functional connectivity related to creativity (Belden et al, 2020; Takeuchi et al, 2010), likely drawing on its role in memory retrieval for the generative and evaluative processes involved in creative behavior. Older adults have also been shown to have higher relative activation than younger adults in LG when comparing positive relative to negative visual stimuli (Kehoe et al, 2013).

Finally, there was a pronounced effect across all subjects, in which listening to their own self-selected music led to increased connectivity between reward network and auditory regions as compared to other recognizable music within the larger western canon. Agency in music listening is known to be an important factor in health outcomes of music-based interventions (Ruud, 1997; Cassidy & Mcdonald, 2009; Howlin et al, 2022). However, here for the first time, we present direct evidence that agency in song selection leads to increased connectivity of auditory and reward networks. This supports the use of these networks as neural targets for music-based interventions that center around the concept of agency, and helps to explain these previously observed behavioral effects.These results constitute a novel set of findings characterizing age-related differences in the musical reward response. These findings suggest that auditory and reward systems can be directly targeted by music-based interventions for older adults, and provide neurological evidence supporting the use of participant-selected music in such interventions.

Previous work has noted the importance of agency for music-based interventions for dementia (Baird & Thompson, 2018). Specifically, the arguments for the sense of agency in music listening have focused on “embodied selfhood” (Kontos, 2014), in that the persistence of musical intention despite advanced Alzheimer’s disease is explained by “corporeality as a source of agency” (Kontos & Martin, 2013): i.e. the predictive processes by which music engages the cognitive system confers cognitive benefit in dementia by restoring a sense of agency. The present study reconciles this idea with the notion of network dedifferentiation in neuroscientific studies of aging (Koen et al., 2020). Specifically, these results showed that for older adults compared to younger adults, music listening engages lower within-network connectivity, but greater out-of-network connectivity between auditory-seeded and reward- seeded networks and out-of-network regions. This suggests that persistent music listening in older adults, such as through receptive music-based interventions (Quinci et al, 2022), may confer an advantage by activating more disparate regions throughout different networks in the brain, especially during self-selected music listening. This is consistent with the de-emphasis of perceptual networks, and the relative upregulation of out-of-network processing, during music listening in older adulthood. Designs of music-based interventions may benefit from this finding by emphasizing strategies that are known to pull in out-of-network processes, such as episodic memory, self-referential processing,and/or “embodied” processing, that are uniquely engaged during music listening, especially self-selected music listening, in older adulthood.

Despite the strengths of the current study in reconciling theoretical and empirical findings across disciplines, some limitations should be addressed. Gender disparities do exist between our two samples: the young adult sample had more female participants than the older adults sample, which was relatively gender-balanced. To account for these differences, we ran follow-up analyses to observe any main effects of gender. No significant effects were observed, suggesting that the gender differences were not confounding the patterns of results. The age span was also not consistent between our two samples. Our older adult sample spanned a total of 35 years of age, while our young adult sample only spanned 5. Follow-up analyses revealed no main effects of within-sample age, and including within-sample age as a covariate did not have a significant impact on the observed effects, thus increasing the confidence with which we claim that effects of age primarily arose from differences between the young and older adult samples.

Our data are also limited in the extent to which participants were asked to report on their feelings about a particular piece. We did not ask about the particular emotions evoked by each piece, nor did we ask participants about the autobiographical context behind their self-selected pieces. In future studies, a more granular approach to emotional classification of pieces may be warranted to better understand the relevant contributions to auditory-reward connectivity.

Furthermore, the young adult sample falls into an age range in which the brain is still developing (Giedd et al, 1999; Sowell et al, 2001), and thus for future analyses we would recommend expanding the sample to include a broader range of the lifespan, covering all age groups.

Nevertheless, this novel set of findings demonstrates that different age groups do in fact vary in their auditory reward processing, further explaining observed differences in musical reward experience (Belfi et al, 2020), and adding to the depth of knowledge required for the understanding of musical reward processing.

## Data availability statement

Preregistration can be found at https://osf.io/zxd42. Statistical maps of all significant results can be found at https://neurovault.org/collections/QSMRNLRW/

## Author contribution

Formal Analysis, Data Curation, Methodology, Software, Validation, Investigation, Visualization, Writing - Original Draft Preparation, Writing - Review & Editing: A.B. Investigation, Project Administration, Visualization, Writing - Original Draft Preparation, Writing - Review & Editing, M.A.Q. Writing - Review & Editing: M.G., N.D., S.B.H. Conceptualization, Data Curation, Funding Acquisition, Investigation, Resources, Supervision, Visualization, Writing - Original Draft Preparation, Writing - Review & Editing: PL.

## Acknowledgements and Funding Information

We acknowledge funding support from NIH R01AG078376, NIH R21AG075232, NSF-BCS 2240330 and NSF-CAREER 1945436 to PL. We acknowledge MIND Lab members Nicholas Kathios, Valerie Goutama, Dayang Gong, Jacob Ostapenko, Grace Neale, Catherine Zhou, Ritu Amarnani, Amira Toivoinen, Itamar Zik, Nicole Page, Jasper Olson, and Parker Tichko for assistance with fMRI data acquisition and analyses. We thank all our participants, and staff of Northeastern University Biomedical Imaging Center (Fred Bidmead, Valur Olafsson, Susan Whitfield-Gabrieli) for help with MRI data acquisition and analysis.

## Supplementary Figures

**Supplementary Figure 1:**
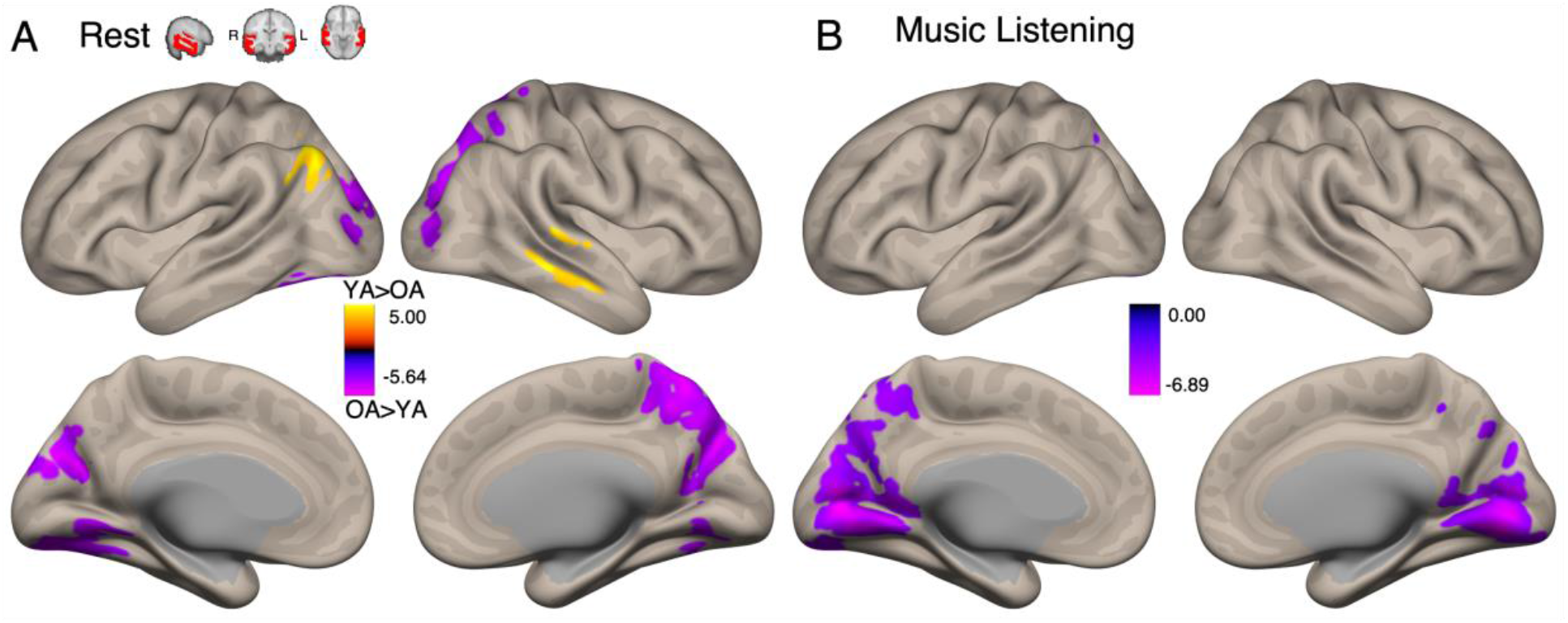
Age Differences in Auditory Network Seed-Based Connectivity Corrected for Differences in BMRQ: Significant clusters for YA>OA contrast in auditory-network seed when including BMRQ as a covariate of no interest, significant at the p < 0.05, FDR-corrected level for both voxel height and cluster size. (A) Visualization of significant clusters during resting state scan, including clusters favoring younger adults (warm tones) and clusters favoring older adults (cool tones). (B) Visualization of significant clusters favoring older adults during music listening task.

**Supplementary Figure 2:**
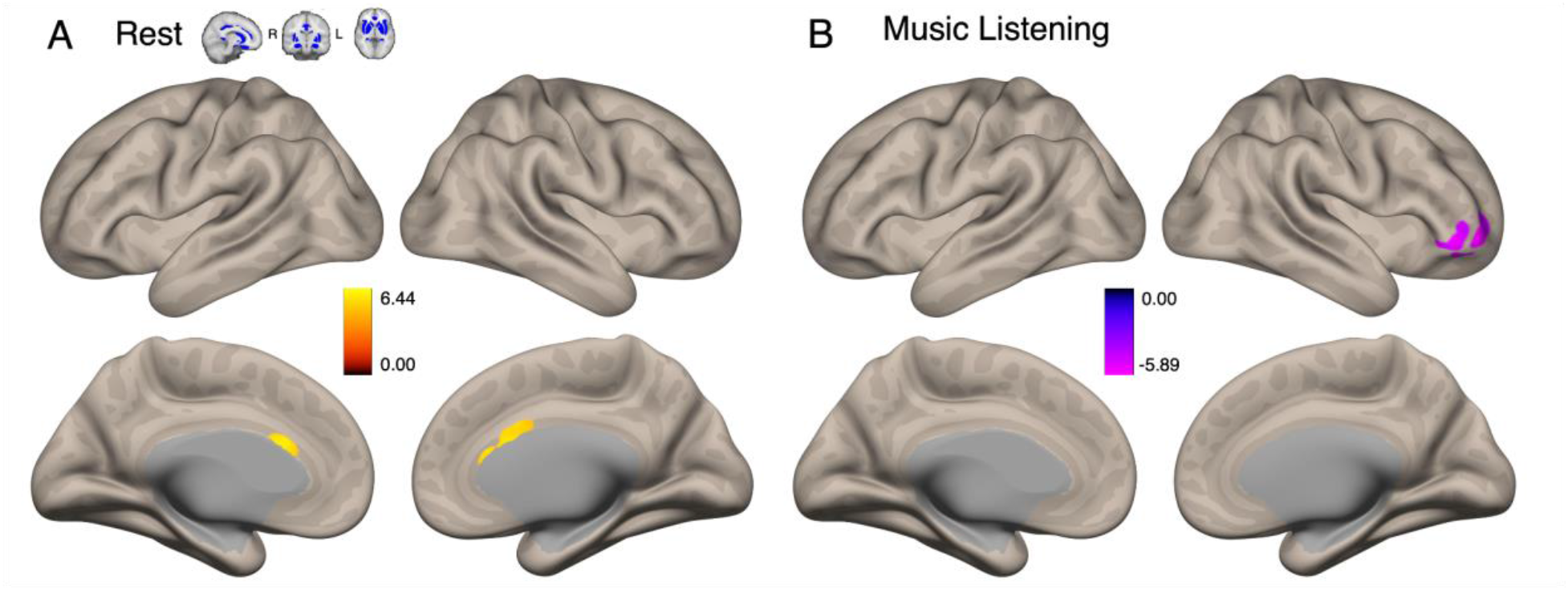
Age Differences in Reward Network Seed-Based Connectivity Corrected for Differences in BMRQ: Significant clusters for YA>OA contrast in reward-network seed when including BMRQ as a covariate of no interest, significant at the p < 0.05, FDR-corrected level for both voxel height and cluster size. (A) Visualization of significant clusters favoring younger adults during resting state scan. (B) Visualization of significant clusters favoring older adults during music listening task.

**Supplementary Figure 3:**
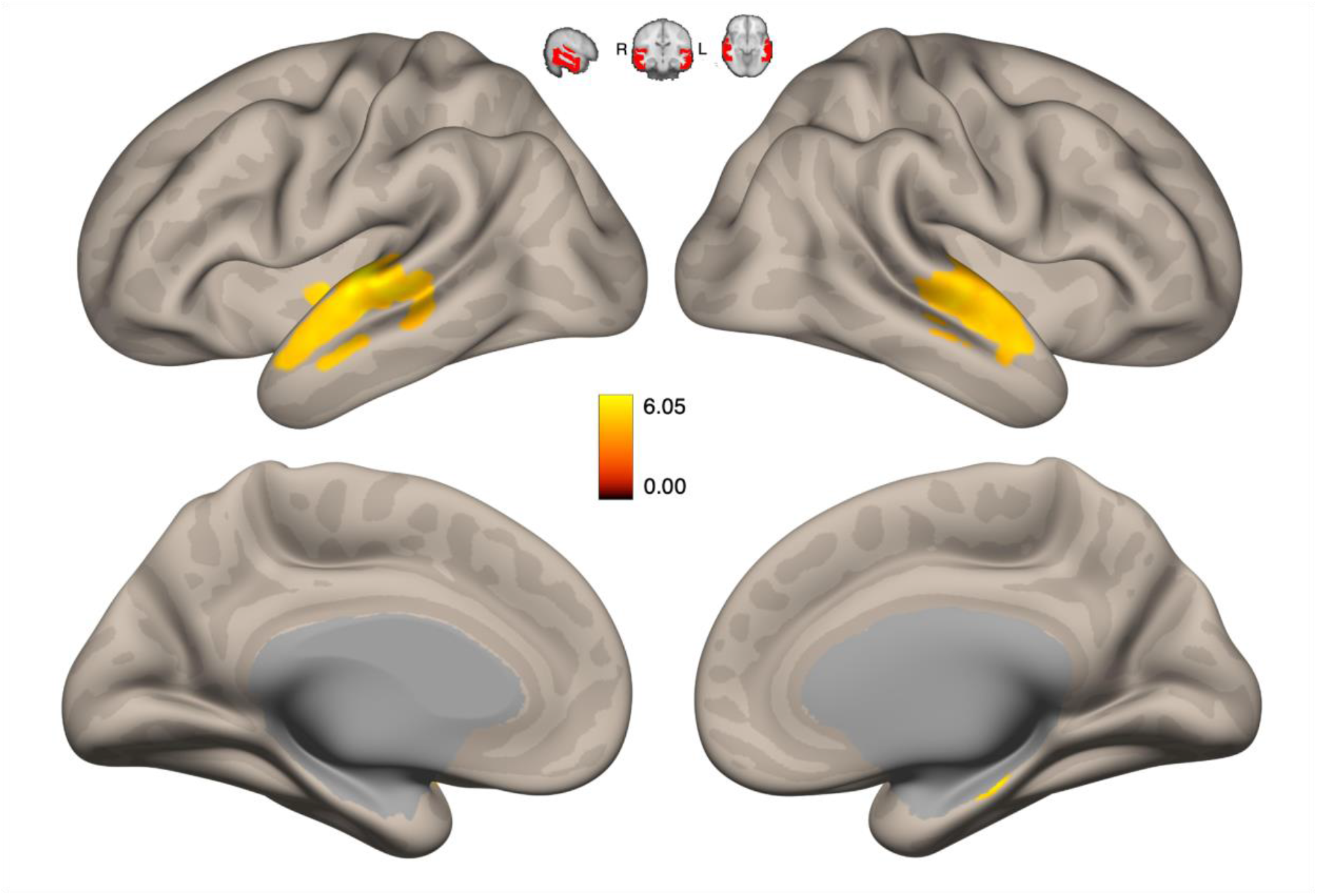
Auditory Network Seed-Based Connectivity of Musical Preference Corrected for Differences in BMRQ: Significant clusters for linear liking contrast (hate, neutral, like, love [-3,-1,1,3]) in auditory-network seed across all subjects when including BMRQ as a covariate of no interest. Clusters are significant at the p < 0.05, FDR-corrected level for both voxel height and cluster size.

**Supplementary Figure 4:**
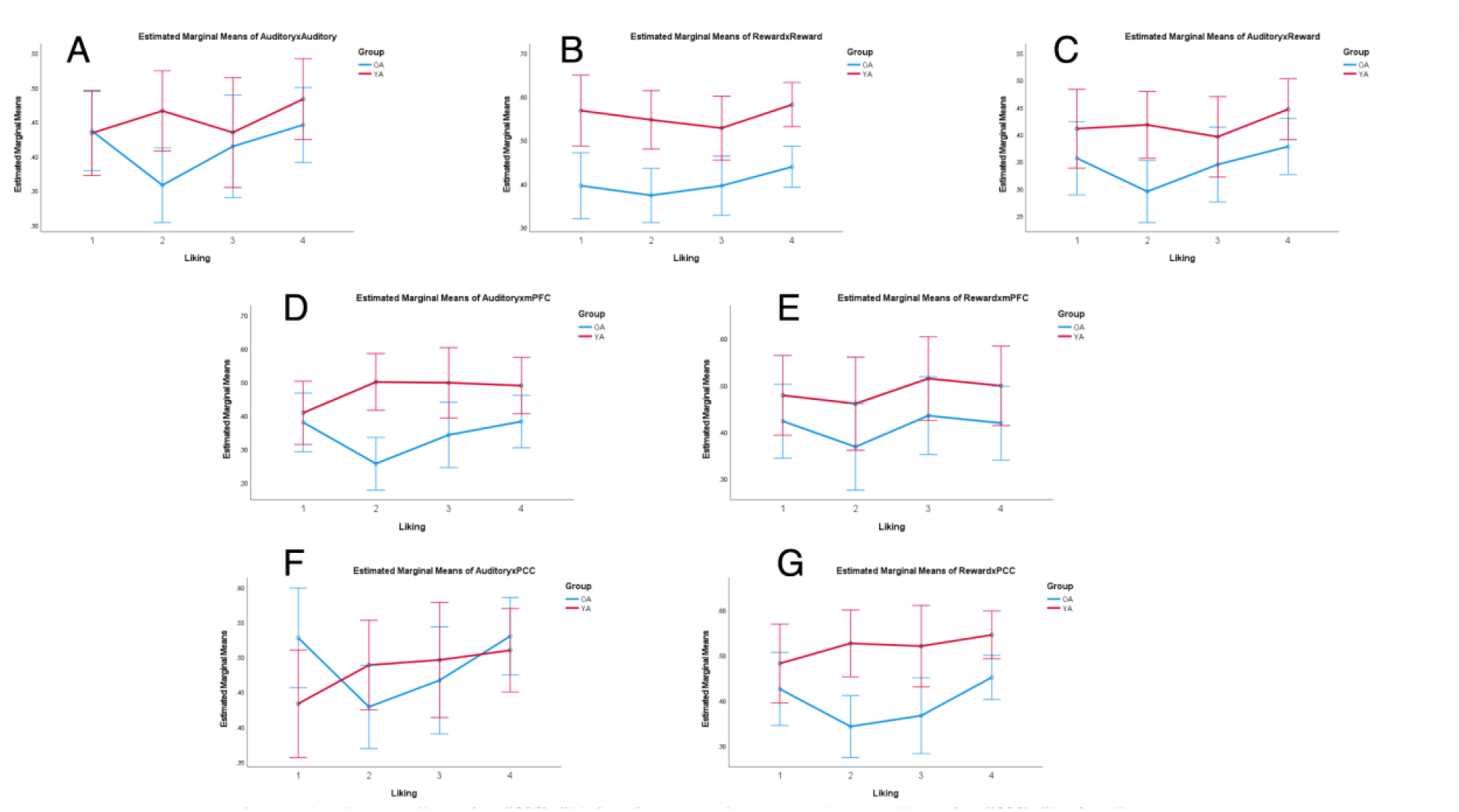
ROI-ROI MANCOVA Results. Bar graph comparing older adults (blue) and younger adults (red) across liking ratings in measures of auditory-auditory (A), reward-reward (B), auditory-reward (C), auditory-mPFC (D), Reward-mPFC (E), Auditory-PCC (F) and reward-PCC (G) connectivity. Error bars represent +/- two standard errors, and covariates appearing in the model are evaluated at the following values: Gender = 0.3947, BMRQ = - 0.3344, within-group age = -0.1491.

